# Closure in the Visual Cortex: How do we sample?

**DOI:** 10.1101/2023.10.20.563146

**Authors:** Irfa Nisar, James H. Elder

**Affiliations:** Department of Electrical Engineering and Computer Science, Center for Vision Research, York University

**Keywords:** shape adaptation, cortical adaptation

## Abstract

Does the human visual system sample shapes at discrete points? During adaptation, when the neurons are fatigued, one observes the underlying principles that were once less prominent than the fatigued features. Operating under deficit, these less prominent features expose the original contributions from the fatigued neurons that are now absent. An underlying lower-order neural process is thus, now revealed. In this paper, we conduct experiments using a modified version of the circle-polygon illusion to reveal the brain’s sampling pattern. The circle-polygon illusion produces polygonal percepts during adaptation when a static dark outline circle is pulsed at 2 Hz alternating with a gradient luminance circle. We define sampling as the edge length of the emergent polygon. We develop a reconstruction function that defines the edge length based on psychophysical responses. We perform two experiments. In the first experiment, we present circles of size [2,4,8,16] deg presented at eccentricity [0,1,2,4,8] deg in a cross design. In the second experiment, we modify the method of Sakurai (2014) and display arc lengths that are 1/8, 1/4, 3/8, 1/2, 5/8, 3/4, 7/8 and 1 (whole) of a circle, of size 4 and 8 deg, presented centrally. The observers report the edge length. We find that the stimulus size and presentation eccentricity, taken together, best explain the edge length reported by the users. The users, as a random effect, do not influence the mean of the edge length reported when considering the best model reported (size and eccentricity together). However, the users do influence edge length reported only when using mean eccentricity or eccentricity as the parameter influencing edge length. Arc lengths of a circle produce the same or similar edge lengths. The length of the curve does not play a significant role signifying that biological neurophysiology at an eccentricity controls the edge length formation. Using the influences on edge length, we define sampling as a sum of qualitative influences and a sampling function derived from Taylors polynomial using sampling values along the eccentricity grid. As we use the sampled values directly to reconstruct the function, we remove the need for recording directly from neurons and instead rely on behavioural responses to build the reconstruction function.

## Introduction

Adaptation is a process by which a set of population neurons are fatigued, allowing other sets of neurons that encode inhibition and are not fatigued, to reveal their neural preference for structure and shape (Anstis, 2013; Ito, 2012). Indirectly, adaptation reveals the original coding that is now absent. In an earlier experiment (Nisar & H. Elder, 2021), regular polygons of varying order were observed during in-place adaptation when a flashing circle alternated with its black outline at 2Hz. We define sampling as the edge length selected in an emergent polygon. In this paper, we attempt to explain how the edge length forms, isolate the cortex region that is engaged in the construction of edges, and reconstruct the function that produces the edge length solely from sampled data values. As we are limited by behavioural data that operates as an aggregated response from biological receptive fields (Heitger, Rosenthaler, Von Der Heydt, Peterhans, & Kübler, 1992; Van Essen, Newsome, & Maunsell, 2002; Pasupathy & Connor, 1999, 2001, 2002), we are unable to probe the individual receptive fields at the psychophysics level. In the absence of linearly separable Gaussian functions derived from neural activations that respond to carefully constructed stimuli (Pasupathy & Connor, 1999, 2001, 2002), we build two experiments to determine what factors influence the edge length and reconstruct the sampling function from sampled data values. In this study, we alter the stimuli, probe if the arc length of the stimulus affects the edge length seen and reveal sampling patterns for curvature closure. We consider the importance of connectedness and, therefore, use arc lengths as the probing parameter. Unlike in (Nisar & H. Elder, 2021), we report edge lengths as it forms the basic precursors to curves through the combination of oriented receptive fields (Orban, 2008). The higher part of the ventral system, region V4 of the cortex, encodes higher-order contour derivatives in the form of tuning for broad curvature as well as certain sharp narrow curvature.Both the contour type and its angular orientation are important descriptors for shape selectivity (Pasupathy & Connor, 1999, 2001; El-Shamayleh & Pasupathy, 2016). (El-Shamayleh & Pasupathy, 2016) show the existence of normalized curvature supported by neurons that fire such that the overall angle subtended by the contour at the center remains the same. However, little has been done to vigorously study the simplest convex shape, the circle, and arc lengths that are part of its circumference, under adaptation, and reveal the encoding at lower levels of the visual cortex. The angle-preserving coding for curvature in mid-level vision might exist earlier in the visual cortex through a different phenomenon. The organization of receptive fields within the cortical column is known to have long-range connections that complete contours (Das & Gilbert, 1997). This may contiguously join discrete edge lengths of fixed orders at a specified eccentricity. We hypothesize that the brain may sample in a predictable way. We know that the sampling is a by-product of closure (MacLeod & Ellis, 1939; Wertheimer, 2007), Wertheimer(1912,1923) from Gestalt theory and also a neuro-physiologically mapping of the arrangement of the receptive fields that may give rise to emergent contours (Gilbert, 1992; Li, Piëch, & Gilbert, 2008). In (Nisar & H. Elder, 2021), in-place adaptation polygons were observed from flashing circles. They were reported to be symmetrical regular polygons, changing strength and polygon order with changing presentation eccentricity and stimulus size. The strength was strongest when the circle was of size 2 degree visual angle and was presented at an eccentricity 8 degree visual angle. The largest polygon order was observed when the percept was of size 16 degree and at an eccentricity 4 degree. The variation in polygon order showed that the lower-level account of human vision was scale-variant. We want to test if the brain codes for edge length at a particular eccentricity in fixed lengths. One way to test this is by varying the arc lengths progressively. We construct two sets of experiments. The first experiment presents full circles with diameter [2,4,8,16] ^°^ and at eccentricity [0,1,2,4,8] ^°^ in a cross design. The second experiment presents arc lengths, that are 1/8, 1/4, 3/8, 1/2, 5/8, 3/4, 7/8, and 1 (whole) of the circumference of the circle, of diameter [4,8] respectively, presented at eccentricity 0. We ask the following questions: (a) How do the edge length and strength of percept change with size? (b) How does the edge length and strength of the percept change with eccentricity? (c) How does the edge length and strength of the percept change with size and eccentricity? Can we build a linear mixed model to characterize the best-fit model from the responses collected? (f) Do participants affect the models? (g) Does M-scaling play a role in predicting the edge length reported? (g) Does the edge length change with changing arc lengths when the circle from which they are generated remains the same? (h) Can we formulate a sampling function on the grid? If the arc lengths do not influence the edge length, we can re-construct the sampling function along the eccentricity grid using the derivatives of the function that are in turn constructed simply from the sampled values on the grid. The key motivation for recording edge length was to identify the unit element that would define the polygon order or shape. We also wanted to eliminate all other theories that would suggest that the edge length was being constructed outside the cortex. We discuss these theories next. Homomorphic class differences are processed earlier and outside the ventral stream. This is revealed in several psycho-physical experiments by Chen and Palmer (Chen, 1982; Palmer & Rock, 1994) . Chen states that connectedness, closed or open, and the number of holes are topological features that form important perceptual representations that are extracted early in the visual process (Chen, 1982). Palmer in (Palmer & Rock, 1994) show that uniform connectedness is a property where regions of similar lightness, colour, or texture are perceived as single units even before grouping principles such as proximity and similarity begin to take effect. Edge length, also defined as the gap between contours, allows the construction of closed shapes if they are small (Elder & Zucker, 1993). This property is called closure. Closure is often thought to be a primitive property, separate from the ventral stream pathway that culminates in the inferotemporal region (IT) that makes inferences about global shape from earlier components. Wertheimer’s earliest experiments in psychology (Wertheimer, 2007) assert that the quality of the part is a function of its integration into the whole. In other words, the whole is more significant and determines the quality of the part. The paradigm of global or local shape are competing features in defining shape (Kimchi, 1992). We, therefore, study edge formation from arc lengths as a corollary to the polygons reported (Nisar & H. Elder, 2021) to see if in-place adaptation to part shapes are different from full circular shapes. If the same edge length is reported regardless of arc lengths, not only can the re-constructed function be applied to any arc stimulus on the grid, but the function can also be approximated to a higher degree of accuracy by including samples from more points along the grid. We can also be confident that the function has a domain in a certain region of the cortex such as region V1.

## Methods

### Observers

30 observers participated in the experiment. All observers participated in the online version of the experiment, hosted on Pavlovia. Participants were recruited using Prolific, an online recruitment site. The participants were screened for being fluent in English and for having corrected-to-normal vision.

### Apparatus

Observers were asked to have a measuring tape for the experiment or discontinue if they did not. They were asked to confirm that they had a monitor screen of atleast 13 inches (or 33 cm) wide and to adjust the chair such that the screen was at a comfortable viewing distance.

To ensure that the correct size and eccentricity of the stimuli was being generated, a set of instructions were displayed at the start of the experiment. The participant reported the viewing distance by measuring the distance (in centimeters) between the bridge of their nose and the center of the screen. The participants were asked to scale a virtual credit card on their screen to fit a physical credit card that they placed against the screen. Pixels per degree was computed by using the scaled card information (in pixels) and the distance of the participant from the monitor.

The size of the stimuli, in degrees and its eccentric placement, was computed and drawn using the pixels per degree information. Other components of the instruction included asking the participant to dim the lights. The background of the experiment was white. The mean luminance of the monitor was expected to read 26*cd/m*^2^.

### Measurement Techniques

Participants were instructed to provide their viewing distance in centimeters (Figure 1(a)) using a measuring tape and scale a credit card to match a physical credit card (Figure 1(b)). The participants indicated the polygon to which they adapted by selecting the edge length in an array of edge lengths during the response selection phase. A standard mouse was assumed to be used to point and click on the edge length since the participants were online. Unlimited time was awarded to select an edge length. Once the edge length was selected, a Likert scale from 1 to 10 was shown to mark the strength of the percept if the participants had indicated a somewhat polygonal percept. Once the participant recorded a response, the next stimuli presentation appeared.

**Figure 1:**
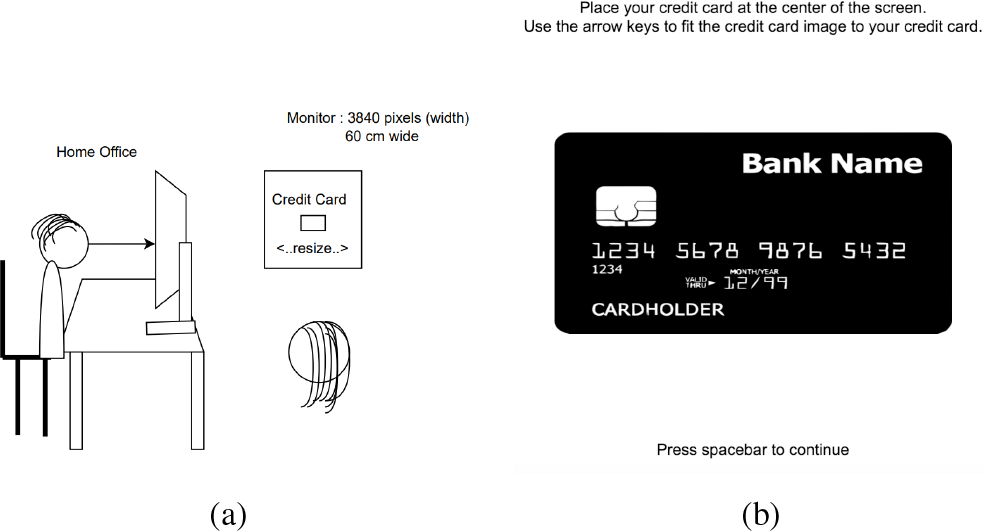
Experiments 1 and 2: Apparatus setup (a) Observer is seated at a comfortable distance from the screen with their head securely held by a chin rest. Observer provides their distance from the screen which helps compute the stimulus size in visual angle degree. (b) Participants were asked to scale the virtual credit card using the arrow keys, such that it matches the length and width of a standard credit card that is placed against the screen. A standard credit card is 8.5 cm wide and 5.4 cm high.

## Experiment 1

The experiment reports the edge length of the polygon percept seen as the circle diameter and eccentricity of the circles placement varies. We present stimuli that are circles and alternate with a circle that is its inward gradation pattern. The diameter of the circles we present as stimuli, subtend a visual angle of 2,4,8 and 16 ^°^ on the users eye. The eccentricity at which the circle center is presented is at 0,1,2,4 and 8 ^°^. There were a total of 36 unique stimuli presentations when accounting for the size, eccentricity and handedness of the stimuli appearance from the fixation cross (left or right). We study the adaptation produced in place, induced by the flashing stimuli, at varying eccentricities along the horizontal meridian and report the resulting polygonal percepts that form. We do not measure the polygon order directly. Instead we use edge lengths that would represent the regular polygon. Participants mark an edge length of the polygon percept they see. The url for the experiment is available at https://run.pavlovia.org/inisar2/fragmentonline30.

### Stimuli

The stimuli consisted of a circle with a black outline that alternated with another circle of the same size but with an inward gradation pattern. The black outlines were constructed of penwidth 5 using Psychtoolbox (Figure 2(b)). The circles were built of different sizes (Figure 2(a)). The eccentricities at which the circle was presented were 0,1,2,4 and 8 ^°^. Randomized trials presented every combination of the circle radius and its eccentricity. Every unique combination of eccentricity and radius was presented ten times for a total of 360 trials. The circle was placed left of the central fixation cross half the time and on the right of the fixation cross half of the time. The stimuli was presented against a white background.

**Figure 2:**
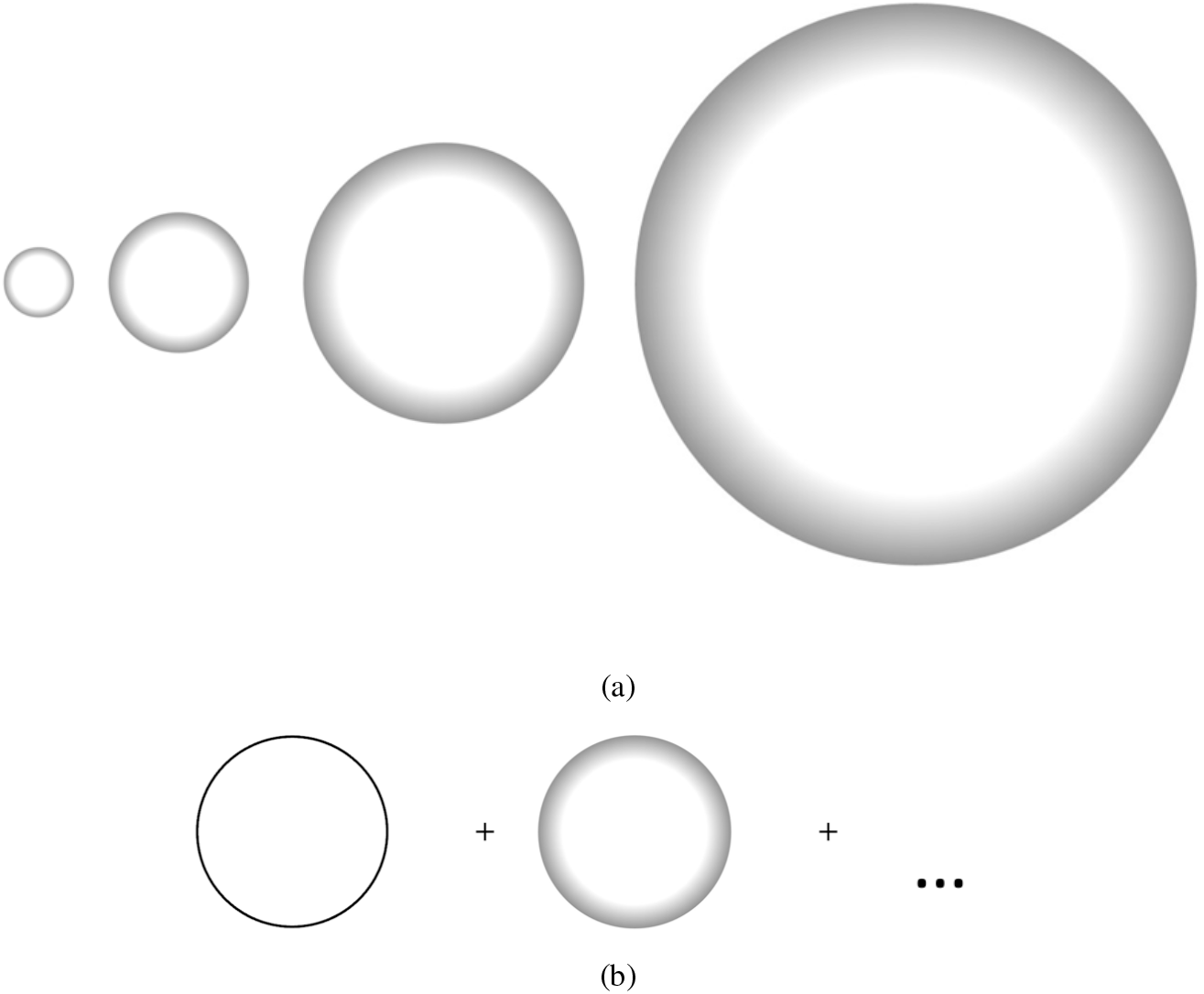
Experiment 1: (a) Stimuli sizes: Circle with inward gradation, whose diameter subtends 2,4,8 and 16 degrees of visual angle respectively on the user’s eye. The pictures are for illustration only and show the relative sizes of the stimuli. Stimuli do not subtend the exact degree on the user’s eye. (b) Flashing stimuli: Circle in a black outline and its inward gradation. The outline and the inward gradation alternate at 2 Hz. The circle is presented to the left of the fixation cross.

## Experiment 2

In this experiment, we construct several circular arc lengths. The arc lengths are 1/8, 1/4, 3/8, 1/2, 5/8, 3/4, 7/8 and 1 (or whole) of a circle. The circle stimuli are of size 4 deg and 8 deg visual degree angle presented centrally. In this experiment we probe the polygon order seen during adaptation using the arc length of the circle. The url for the experiment is available at https://run.pavlovia.org/inisar2/fragmentonline29.

### Stimuli

The stimuli consisted of a circle or an arc of a circle with a black outline that alternated with another circle or arc of a circle of the same size but with an inward gradation pattern. See Figure 3. The stimuli were arc lengths of a circle. The arc lengths were 1/8,1/4,3/8,1/2,5/8,3/4,7/8, and the whole of a circle. Two stimuli sizes were used. One was a size of 4 ^°^ and the other was a size of 8 ^°^. There were a total of 16 unique stimuli when allowing both size and arc length. All stimuli were presented at 0 ^°^ eccentricity. The black outlines were constructed of pen width 5 using Psychtoolbox. The renderings were hosted on Pavlovia. Randomized trials presented each stimulus 10 times. There were a total of 160 trials. Participants were allowed 16 practice trials.

**Figure 3:**
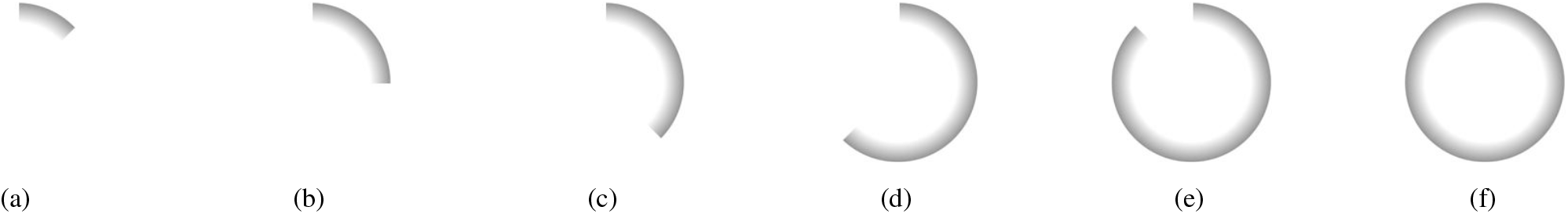
Experiment 2: Graded luminance circle stimulus alternated with its corresponding black outline (black outlines are not shown for brevity). (a) Arc length that is 1/8 of a circle (b) Arc length that is 1/4 of circle (c) Arc length that is 3/8 of a circle (d) Arc length that is 5/8 of a circle (e) Arc length that is 7/8 of a circle. Half and three-quarters of a circle are left out of the figures for brevity.

**Figure 4:**
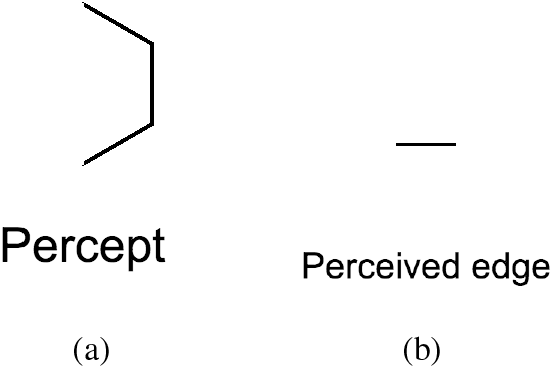
Experiments 1 and 2: (a) Percept shown in the experiment brief. (b) The perceived edge that the participant would have to mark was shown to the participants.

### Procedure

The participants were asked if the stimulus seen was purely circular or somewhat polygonal in an alternative two-forced choice. Circle selections were automatically assigned a strength of 0. If the participants indicated it was somewhat polygonal, they were then asked to judge the perceived edge length and the strength of the percept between 1 to 10. Instructions at the start of the experiment explained to the participants that they would be marking the edge seen in 4(b) if they saw a percept in 4(a). The perceived edge lengths were displayed as a series of twenty line segments built from the equation *log*(*ab*^*c*^) such that a = 1 and b = 0.9. c belonged to the range [1, 20] and each value of c gave rise to a unique line segment (Figure 5 (a)). The difference between the line lengths, when they were represented in their log formats, was uniform. The shortest line segment subtended 0.105 ^°^ visual angle on the eye whereas the largest line segment subtended 2.107 ^°^ visual angle on the users eye.

**Figure 5:**
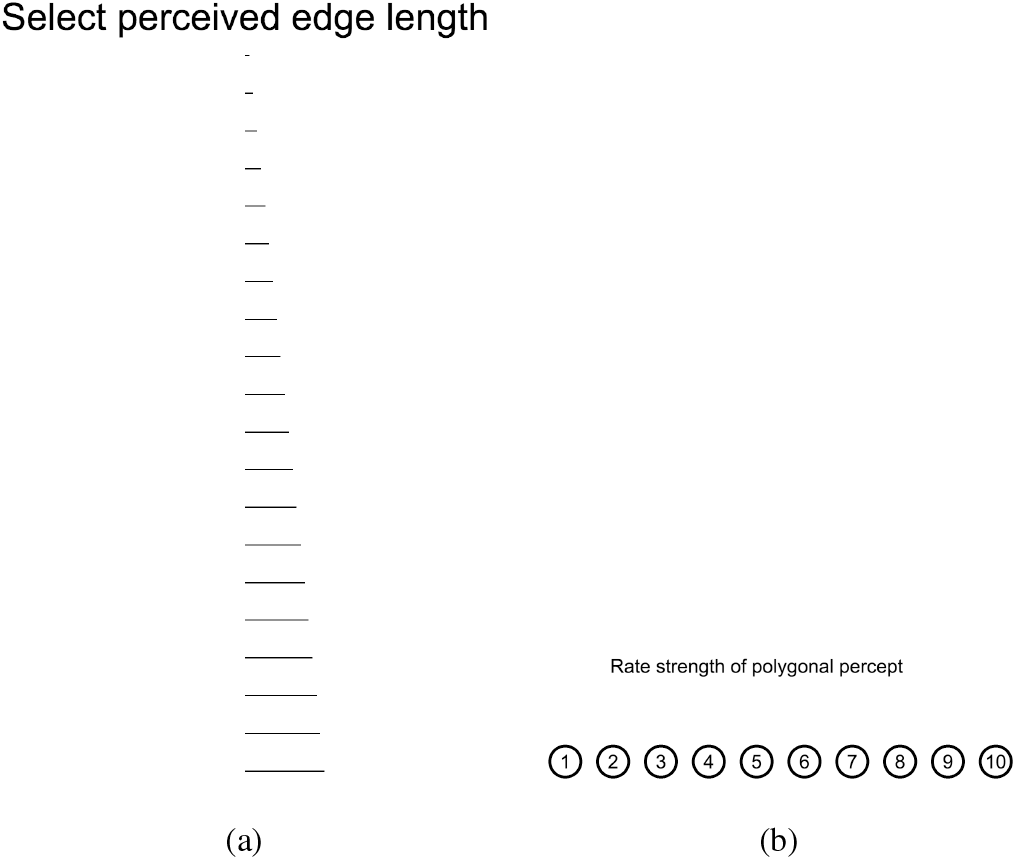
Experiments 1 and 2: (a) Users were asked to select the perceived edge from an array of uniformly incremented log-length lines. (b) Users were asked to rate the strength of the polygon percept from 1-10.

Prior to starting the experiment, the observers were briefed with the following instructions: “After the first phase of the trial, you will be asked if you saw a perfect circle or a polygonal percept. If you perceive the shape as somewhat polygonal, you will indicate the perceived length of the polygonal edges. An example of the polygon seen and the corresponding edge length that you will mark is shown on the right. If you indicated a polygon percept, you will indicate the strength of the polygon percept on a scale from 1 (very weak) to 10 (very strong).” (Figure 5(b))

The participants were asked for their viewing distance from the screen by measuring and reporting the distance (in centimeters) between the bridge of their nose and the center of the screen. To ensure that the stimuli were the right size, the participants were asked to scale an image of a credit card displayed on the screen till it matched a physical credit card using the arrow keys. The scaling allowed the computation of pixels per degree for the participants monitor screen and the appropriate display of the stimuli in degrees thereafter.

“

### Results from Experiment 1

#### Shape

42 percent of the 30 observers reported seeing polygons (Figure 6(a)). As the eccentricity increases, the proportion of polygons seen increases (Figure 6(b)). We see in Figure 10(b,c) that the proportion of polygons reported and the strength of seeing one are not co-related. An observer either sees a polygon or does not but when a polygon is reported, the strength of that polygon is reasonably high. Thus, we can use the averaged values for reconstructing sampling functions on the cortex. Retinotopical organization in human vision is designed such that as the density of the lateral ganglion cells declines in peripheral vision, the receptive fields become larger (Rovamo & Virsu, 1979). Closer to the fovea, the lateral ganglion cells are denser and have smaller receptive fields. The density of nerves at the retina feeds a fixed span of cortex and the human magnification factor, M, is related to the eccentricity of presentation as an inverted cubic equation. As the stimulus moves from the fovea to the periphery keeping the subtended retinal dimension the same, the cortical image remains unchanged because the retinal stimulus is scaled by M onto the cortex. In a similar vein, Harvey et. al (Harvey & Dumoulin, 2011) also show that the regions of the cortex sample lower regions of the cortex such that as the cortical magnification decreases, the receptive field size increases, thereby preserving the point image through the cortex. However, despite the idea of a point image scaling by a cortical magnification on the visual cortex such that the perception of the overall retinal magnification remains the same, our experiment shows that the amount of shape and polygon reported changes with eccentricity. This has consequences for the scaling invariance property which we refute in this series of experiments. We will also show in this section that region V1 of the cortex is the most probable region where the edges are formed during in-place adaptation. A two-sample t-test of proportion polygons at eccentricities 0 and 8 deg shows that the probability of polygons reported have significant mean differences (H=1,p*<*0.05). We see significantly more polygons reported at larger eccentricities (Figure 6(b), 6(c)) than at eccentricity 0. This has at least one implication. The presence of a more noticeable edge length occurs in regions with larger receptive fields such as the periphery. Even though the resolution of the retinal stimulus is expected to be preserved on the cortex, the circle-polygon phenomenon producing edges after adaptation is not guaranteed. Closer to the fovea, receptive fields at 0 ^°^ eccentricity show the least amount of emergent edges.

**Figure 6:**
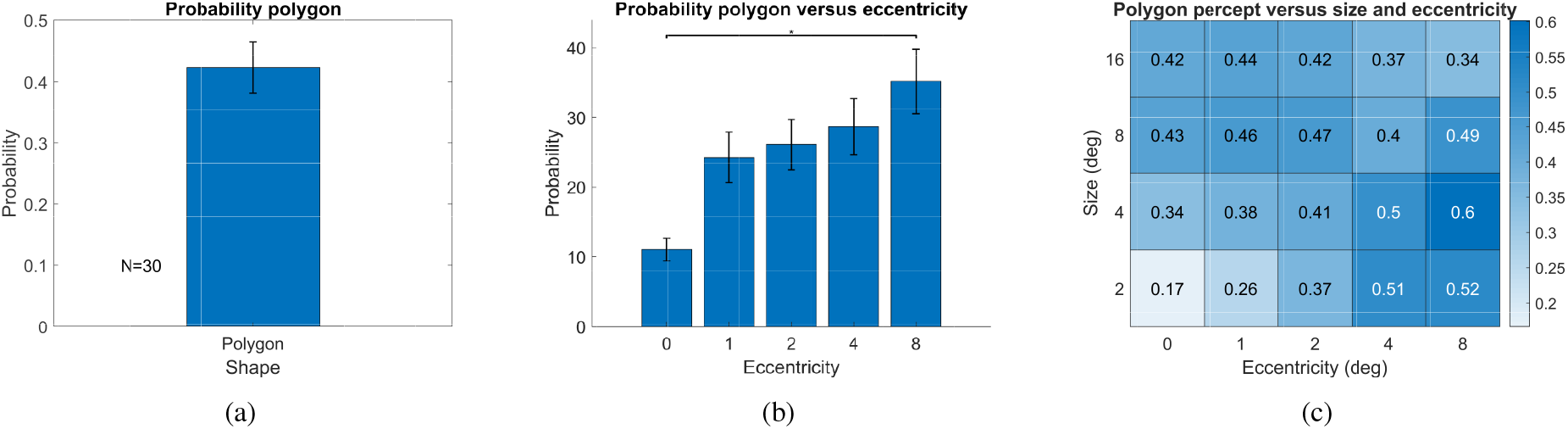
Experiment 1: Probability of seeing polygons versus circles. (a) Probability of seeing polygons over all eccentricities and all stimulus sizes. (b) Probability of seeing polygons at eccentricities 0,1,2,4 and 8 deg visual angles. (c) Probability of seeing polygons at sizes 2,4,8 and 16 deg and eccentricities 0,1,2,4 and 8 deg. Error bars are standard errors of the mean.

#### Edge Length

Edge length increases with both size and eccentricity (Figure 7(a,b), 8(a)). Size is a larger effect and eccentricity is a marginal effect (size:F=80.23,p*<*0.05,*η*^2^ = 0.0215; eccentricity:F=31.39,p*<*0.05,*η*^2^ = 0.0114).

**Figure 7:**
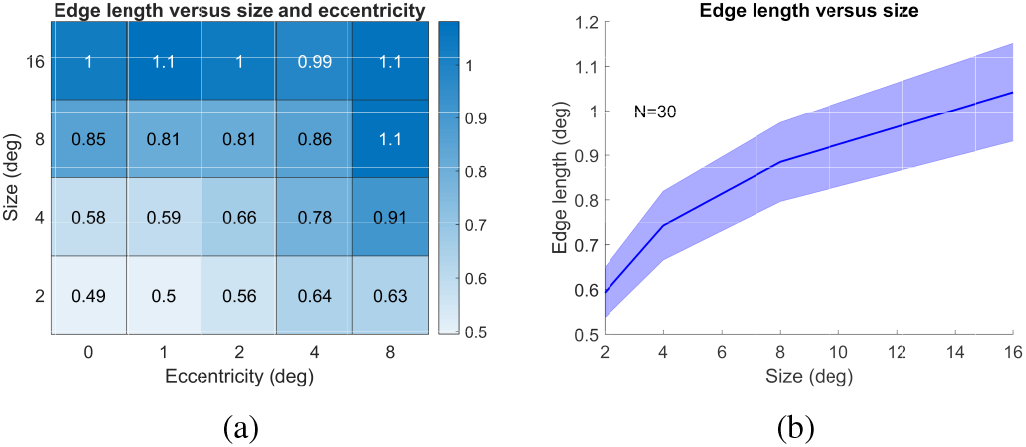
(a) Mean edge length over all sizes of stimulus and eccentricities (b) Mean edge length versus size. Shaded bars are standard error of the mean centred around the mean.

The stimuli is presented at a given eccentricity and is of a prescribed visual angle degree in size. A stimuli can be quite large and may have peripheral influences from its own shape affecting in-place adaptation. Each point on the circumference of the circle subtends a particular visual angle degree on the participants eye. We define minimum eccentricity as the smallest eccentricity that is subtended on the participants eye from any point along the stimulus circumference (Figure 9(a,b)). The mean eccentricity is the summation of subtended eccentricity along the entire circumference (Equation 1, 9(c)). We find that edge lengths are not better explained by either the minimum or mean eccentricity (Table 1, Figure 8(b,c)). Perhaps, the influences can be rephrased as additive across the receptive field overlaps. If we think of the receptive field as a polar grid with the grid lines drawn along the lines enclosing the edge length, we define a sampling field where the influences are additive. This theory is undertaken in the modeling section.

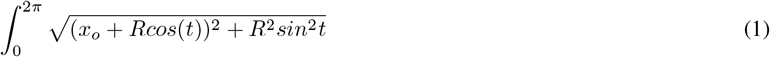

**Figure 8:**
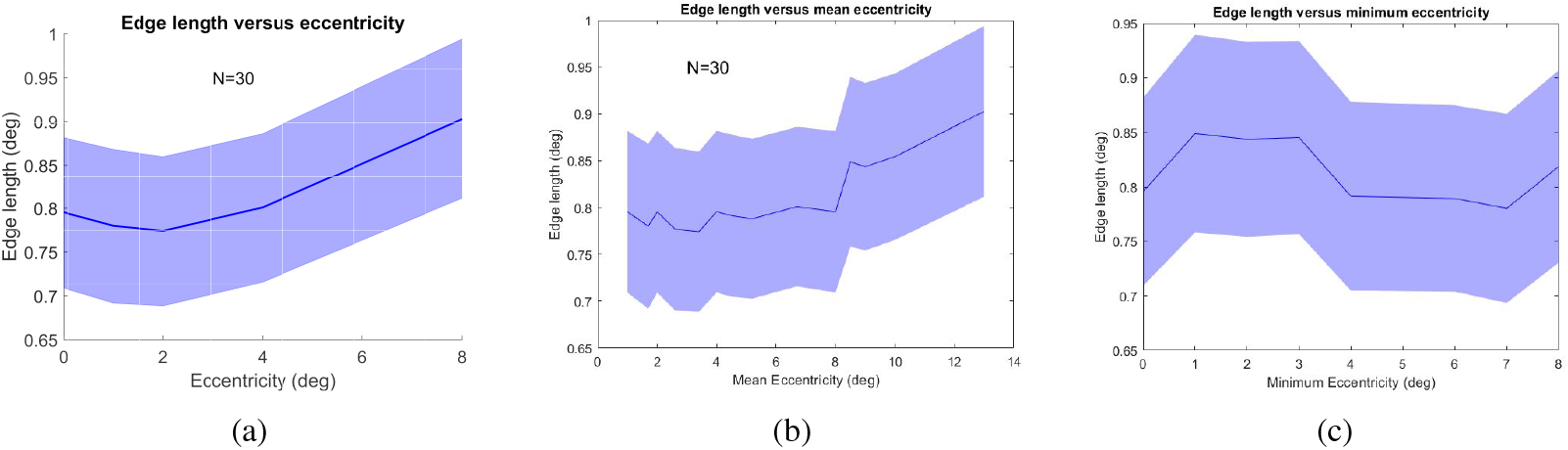
Experiment 1: (a) Edge length versus eccentricity (b) Edge length versus mean eccentricity (c) Edge length versus minimum eccentricity. Edge length increases with eccentricity for regular eccentricity and mean eccentricity but not for minimum eccentricity. Means are computed over all participants. The shaded error bar is the standard error of the mean spread around the central mean line.

**Table 1:**
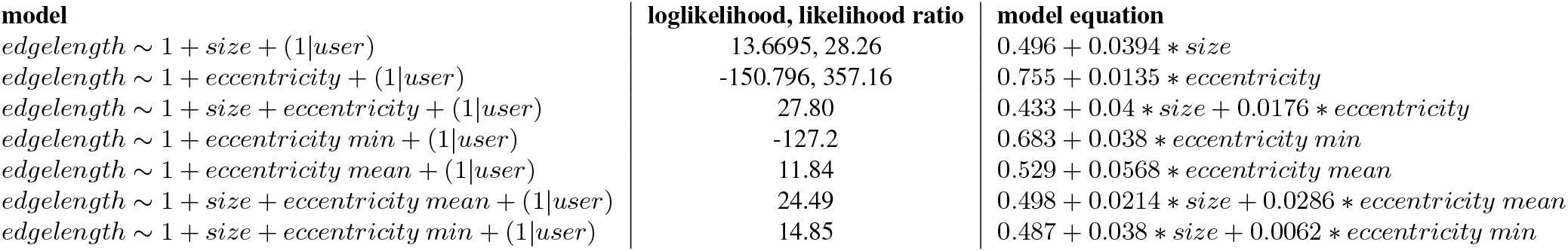
Models and loglikelihood.

**Figure 9:**
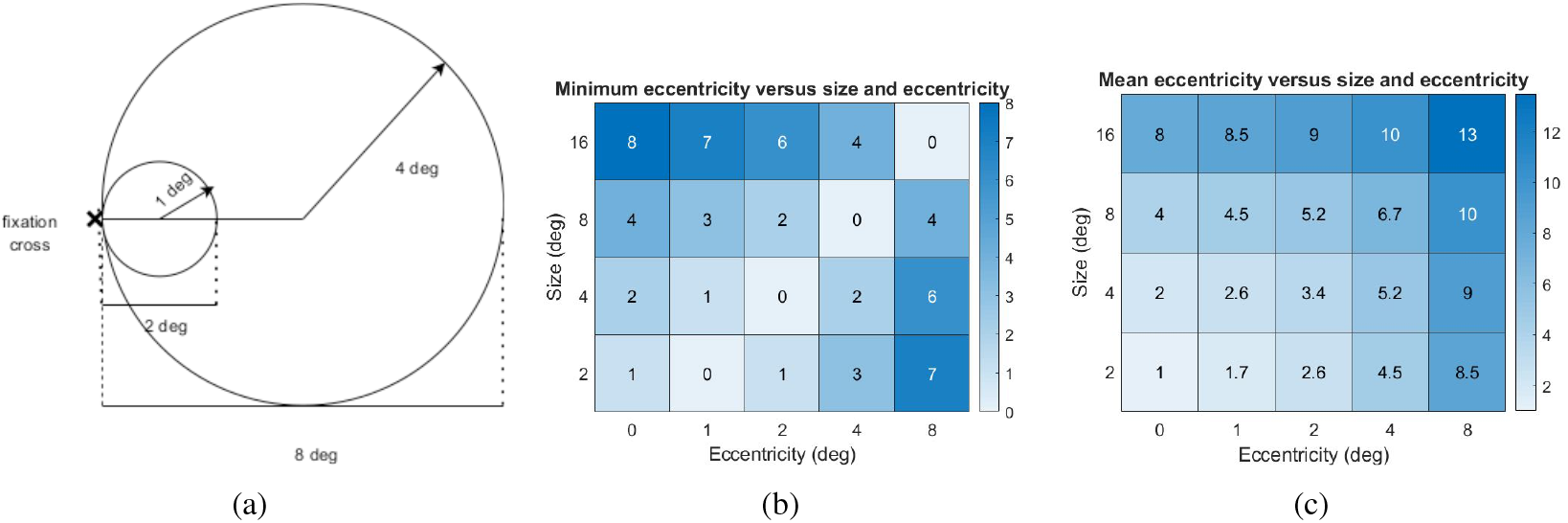
Experiment 1: (a) Both a stimulus of size (diameter) 8 deg and a stimulus of size 2 deg subtend a minimum 0 deg visual angle at the participants eye fixated at the central cross. (b) Minimum eccentricity subtended at the participants eye given the stimulus at size and eccentricity. (c) Mean eccentricity summed along the stimulus circumference and subtended at the participants eye at the given eccentricity.

**Figure 10:**
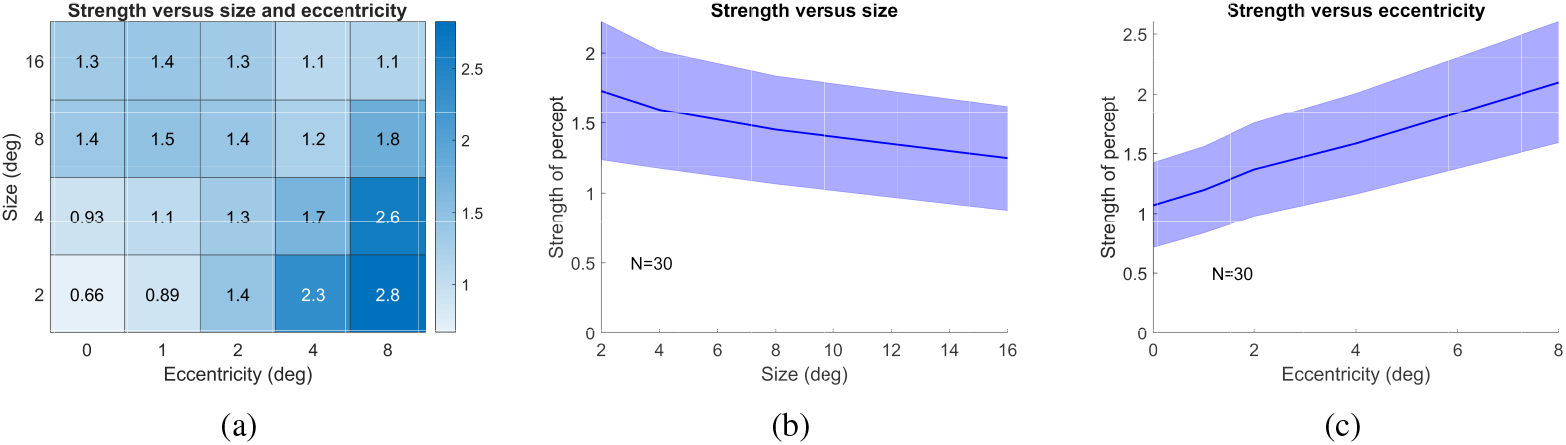
Experiment 1: Strength as a function of size and eccentricity (b) Strength versus eccentricity (c) Strength versus size. Shaded error bar is standard error of the mean centred around the mean.

The edge length is a product of certain neuro-physiological features: (a) the receptive field (Rovamo & Virsu, 1979) (b) the presence of antagonistic receptive fields that may truncate or suppress the formation of an edge in a direction (Heitger et al., 1992; Das & Gilbert, 1997; Orban, 2008) and (c) the contributions from receptive fields that overlap outside the cortical column (Das & Gilbert, 1997). We show that the receptive fields for the edge length we are reporting are located in region V1 of the cortex. Neuro-physiological studies operate at a cellular level allowing direct modelling equations to be built from activated or isolated biological units such as receptive fields or a host of neuron activations. Even so, the equations are often a product of linearly separable Gaussians fitted to activation functions and offer localized equations (Pasupathy & Connor, 1999, 2001, 2002). Psychophysics, on the other hand, is a higher-level behavioural approximation of these activations. In the experiments, we suffer from the same limitations of recording weaker signals captured from behavioural interactions with the stimulus. However, we will use theoretical concepts from neurophysiology to offer a numerical solution to observed biological behaviour.

#### Strength

The experiment measured the strength of the percept on a scale from 1 to 10. 0 meant that the observer did not see a polygon percept after the adaptation. The percept in this case remained the same as the circular stimuli. A 10 on the scale indicated a crisp regular polygon. The strongest effect is for small objects at large eccentricities (Figure 10(a)). Large objects at small eccentricities and small objects at large eccentricities produce large effects. Small objects at small eccentricities and large objects at large eccentricities produce smaller effects (Figure 10(b,c)). Eccentricity is a small effect on the strength reported (*η*^2^ = 0.02, p *<* 0.05) and size is a negligible effect (*η*^2^= 0.004, p *<* 0.05).

We define the sensitivity of positively seeing the stimulus using a sensitivity parameter, z. We define hit as seeing the polygon, a miss as only seeing the circle. We define d prime as a z-transform of polygon hits minus a z-transform of circle cases. d prime is not co-related to the proportional polygons reported as seen from Figure 11(b,c). We show the strength reported and the proportion of times a polygon is reported are not co-related (Figure 11(a,b,c)). When a polygon is reported, the strength is high (Figure 11(a)).

**Figure 11:**
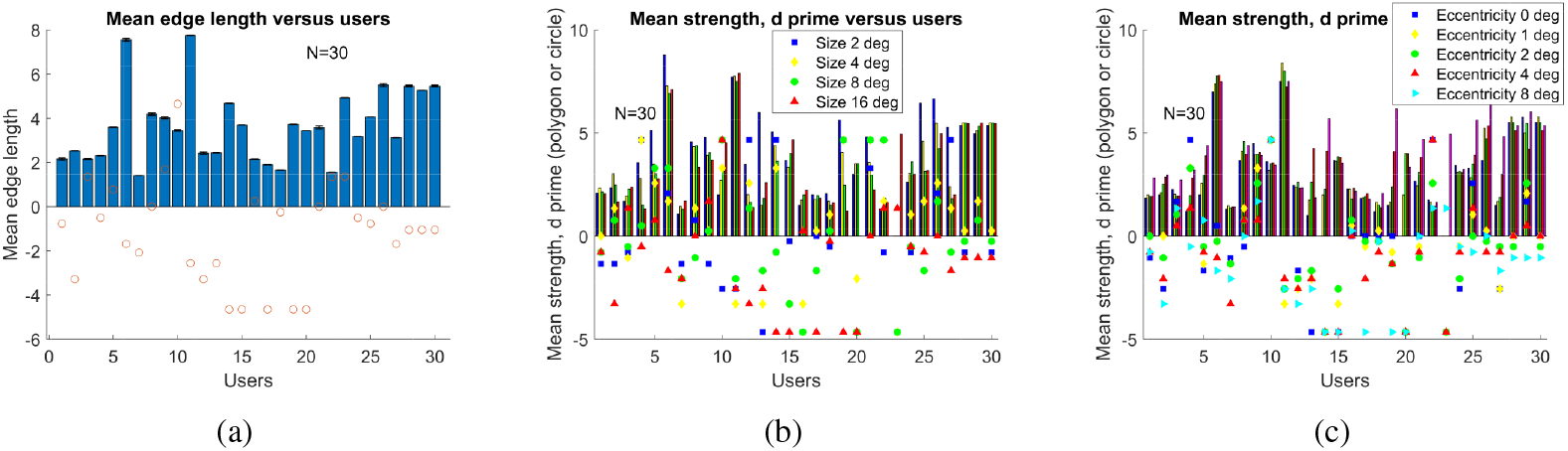
Experiment 1: d prime is the difference of the z-tranform of seeing a polygon and seeing circles. (a) Mean edge length for each user across all size and eccentricity stims. Circles are the d prime reported for the user. (b) Mean strength of polygon reported per user for each size of stimuli. d prime is shown as shaped icons for the various sizes (2: blue square , 4: yellow diamond, 8: green circle , 16:red triangle). (c) Mean strength of polygon reported per user for each eccentricity. d prime is shown as shaped icons for the various eccentricities (0: blue square, 1: yellow diamond , 2: green circle, 4: red triangle , 8: cyan triangle).

#### Linear Model: Identifying factors

The edge length formed during adaptation is a function of several parameters. Receptive fields in cortical columns organize receptive fields of a particular orientation cluster together (Das & Gilbert, 1997) allowing precise activation for oriented edges. Lateral horizontal connections between columns complete or alter an edge length to form contours (Li et al., 2008). An edge length itself is a function of several theories: (a) Orthogonal receptive fields that are antagonistic in nature truncate an emerging edge length (Orban, 2008). Due to two antagonistically aligned orthogonal receptive fields, two orthogonal edges may arise. In such vector space like fields, an edge length operates under an operation like a dot product. In the absence of exact orthogonality, antagonistic orientation may produce edges that enclose angles other than right angles. (b) Due to the clustering nature of similarly directed receptive fields in the cortical columns, the space of operations such as the activation for edges oriented in a particular direction is pre-determined. (c) The stimulus’s size may activate several receptive fields across the visual field and peripheral parts of the stimulus may influence the final emergent edge length at the central fixation cross. In order to understand the main factors contributing to the emergent edge length, we regress edge length across all the factors and study each model for the most probable model and factors. We will use the learnt qualitative factors for building a reconstruction function designed by the nature of contributing factors.

### Results from Experiment 2

Participants did not report twice as large an edge length with twice as large a stimulus (Figure 12(a)). The edge length reported per arc length did not change greatly with arc length but changed with actual circle size (Figure 12(a)). This demonstrates that sampling is based on the quality of the curve or curvature rather than the length of the curve. The strength reported was slightly lower for full circles compared to fractional arc lengths, alluding that the visual system has an affinity for closure and prefers to see shapes as composited closed circles where possible (Figure 12(b)).

**Figure 12:**
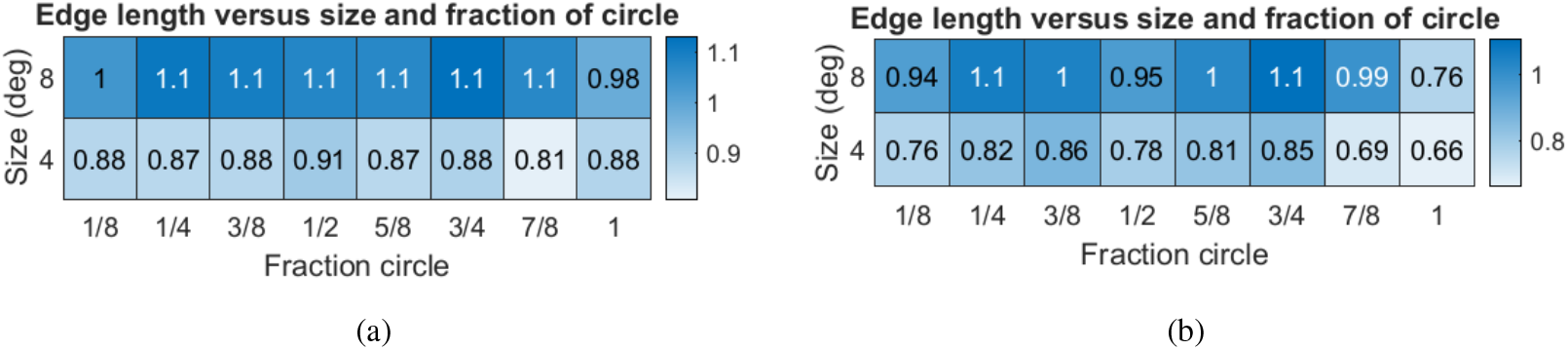
Data from Experiment 2: (a) Mean edge length (in pixels) for different arc lengths 1/8 ,1/4 ,3/8 ,1/2 ,5/8 ,3/4 , and 1 of the circle, circle of deg 4 and 8. (b) Strength reported for different arc lengths 1/8 ,1/4 ,3/8 ,1/2 ,5/8,3/4, and 1 of the circle, circle of deg 4 and 8. Strength was marked on a scale of 1-10 for polygonal percepts. The lowest strength was reported for full circles.

## Discussion

Rovamo in (Rovamo & Virsu, 1979) show that a point on the 2D retinal visual field can be directly scaled and mapped to a specified region on the cortex. The coefficient that maps one degree of visual angle on the retinal plane to a certain milimeter of the cortex is called the linear cortical magnification factor, M. The absence of a flat line when measuring the same parameter using systems neuroscience recordings and behavioural neuroscience reveals differences in the equivalency of the two systems. A granular biological recording of curvature component or features does not necessarily translate to an aggregated response at the behavioural level as reported by participant recordings. Figure 13 (c) demonstrates that behavioural recordings for edge lengths at 0,1,2,4 and 8 ^°^ differ from the M-scale values at the same eccentricities (red line) in a non-linear manner. A ratio of the M scale value over the edge length at eccentricity should be a flat or linear line but is instead a higher order cubic (blue line).

**Figure 13:**
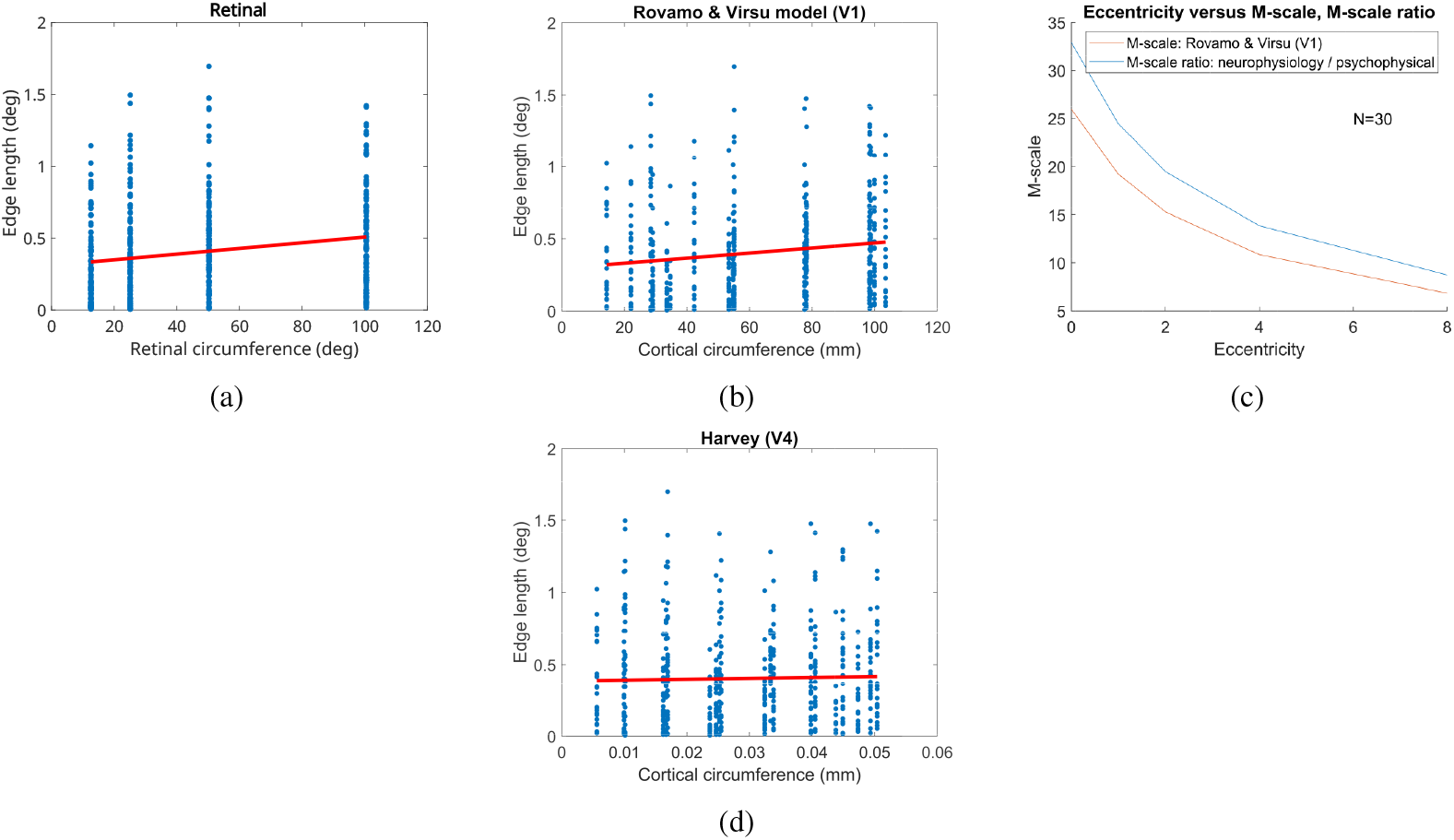
Experiment 1: (a) A bootstrapped linear regression performed to regress edge length as a function of the retinal circumference *R*^2^ = 0.04. (b) A bootstrapped linear regression was performed to regress edge length as a function of the cortical circumference in region V1 *R*^2^ = 0.023. (c) M-scaling shows how a point (unit element) presented at eccentricity 0,1,2,4 and 8 ^°^ scales on the cortex (red line). A ratio of the M-scaling that shows the cortical circumference built from a psychophysical response over retinal circumference scaled by neuro-physiological M-scaling onto the cortex. A ratio of the M-scaling from experimental observer data and neuro-physiological study does not observe a flat slope indicating that animal studies and human data show a linear co-relation but are not equivalent. (d) A bootstrapped linear regression was performed to regress edge length as a function of cortical circumference in region V4 of the cortex *R*^2^ = 0.0007

Next, in 13(a,b,d), using bootstrapped sample and replacement from the set of participant recordings, we regress the edge length against the magnified cortical circumference in regions V1 and V4 (Appendix Algorithm 1) and fit a first-order linear regression model of the edge length as a function of the circumference of the stimuli, magnified on the cortex region and unmagnified on the retina. We find that the edge length is better explained by earlier regions of the cortex. During cortical magnification, we use the M-scale coefficient value for the particular regions of the cortex (Rovamo & Virsu, 1979; Harvey & Dumoulin, 2011). We find the regression *R*^2^ to be higher for region V1 of the cortex than region V4 indicating edge lengths are likely formed in region V1 of the cortex.

### Linear regression model

From Table 1, we fit several models with different predictors to the data and use log likelihood and the likelihood ratio test to find the best fit model. A model with both size and eccentricity allowed to vary freely instead of being fixed at zero, reports the highest log likelihood and forms the best model. With respect to size and eccentricity, we compute the likelihood ratio test to estimate if adding more parameters (size and eccentricity) improves the model with just size or just eccentricity significantly. As indicated by the likelihood ratio test,

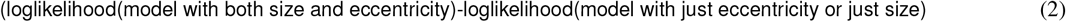

reports a high test statistic 28.26 (size),357.16 (eccentricity) indicating that the model with both predictors (size and eccentricity) fits the model better than the model with just one predictor.

The stimuli is presented at a given eccentricity and for a particular degree visual angle size. The stimuli can be quite large and may have peripheral influences from its own shape effecting in-place adaptation. However, we note that the minimum and maximum eccentricity do not influence edge length as strongly. This shows that peripheral influences do not have a major skewing influence at the central fixation cross.

Cortical magnification would magnify an edge length formed in the cortex region V1 by the following equation in Eqn 3 that relates eccentric presentation to the cortical magnification factor M. *M*_*o*_ is the magnification at the fovea center defined as 7.99 mm/deg.

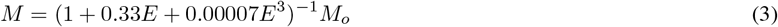

Similarly, it can be argued that the shape represented by the cortex area is predictibly scaled and preserved as sampling also exists through population receptive fields (Harvey & Dumoulin, 2011). However, we find no indication that the cortical magnification factor, M-scale, follows the same scaling pattern as the circumference formed on the cortex. As seen in 13(c), a flat line (blue line) at unity is absent meaning that neurophysiology (M-scale value) at each eccentricity and psychophysical edge length reading at the same eccentricity as a ratio of the M-scale value does not scale linearly. In light of a missing direct relationship, we attempt a sampling function that is purely derived from user reported edge length values.

A Taylor’s polynomial offers an approximation of a function f(x) in a small region around x, *x* + *h*, using the derivatives of f(x).

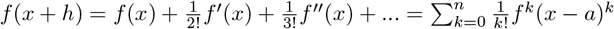

Using forward differencing, we approximate the first and second derivatives of a given function f, as,

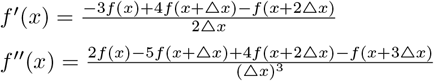

Sampled values at f(x+h),f(x+2h),f(x+3h) are then substituted to approximate f’(x) and f”(x). We obtain an approximation to f(0) using f(x), f’(x) and f”(x) as,

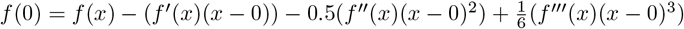

Thus, we are able to reconstruct the original function at f(x), the central fixation cross at f(0). Figure 14 shows the Taylor’s approximation of the reconstructed function at the fixation cross across several size stimulus. The edge length that is approximated follows a similar monotonic increasing pattern as seen in Figure 7(a) at eccentriciy 0 ^°^.

**Figure 14:**
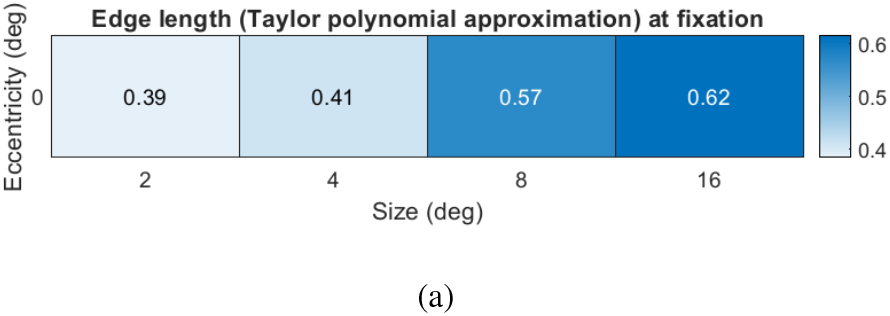
Experiment 1: Taylor’s polynomial approximates the edge length at fixation cross using the sampled values at the grid points 1 ^°^, 2 ^°^, 4 ^°^ and 8 ^°^ using forward differencing to estimate the edge length at the central fixation cross. While the edge length values do not exactly mirror the recorded psychophysical values, they closely follow the values.

We offer this approximation as the reconstructed function for deducing edge length (and therefore early curvature formation). We will only use the sampled values of the function at the grid points to reconstruct the original sampling function. This removes any presumptions of defining a precise function that produces the edge length are constructed in V1.

As we sampled the edge lengths at eccentricities 0,1,2,4, and 8 ^°^, we will use the edge length values at f(x+h), f(x+2h), f(x+4h),f(x+8h) to reconstruct the function at the central fixation cross f(x). If our approximated reconstructed function f(x) matches the edge length reported at the center, we will have obtained a reasonable way to mathematically represent the function. As mentioned in (Rovamo & Virsu, 1979), M is measured along the horizontal meridian and best represents the scaling along the horizontal meridian. As a result, approximating f along the horizontal meridian should provide the closest reconstruction of the function.

## Appendix

We define a simple way to compute curvature on the visual cortex using the tangent at a point. We compute tiny arc lengths at these points using the tangent at the point. We integrate the tiny arcs to obtain the entire curve traced. Since our starting image, the circle, is at the retina, we project points that form the retinal circumference onto the cortex using the human cortical magnification factor, M. M is well defined in (Rovamo Virsu,1979) along the horizontal meridian of the visual field.

### Algorithm 1

A cortical circumference mapping function

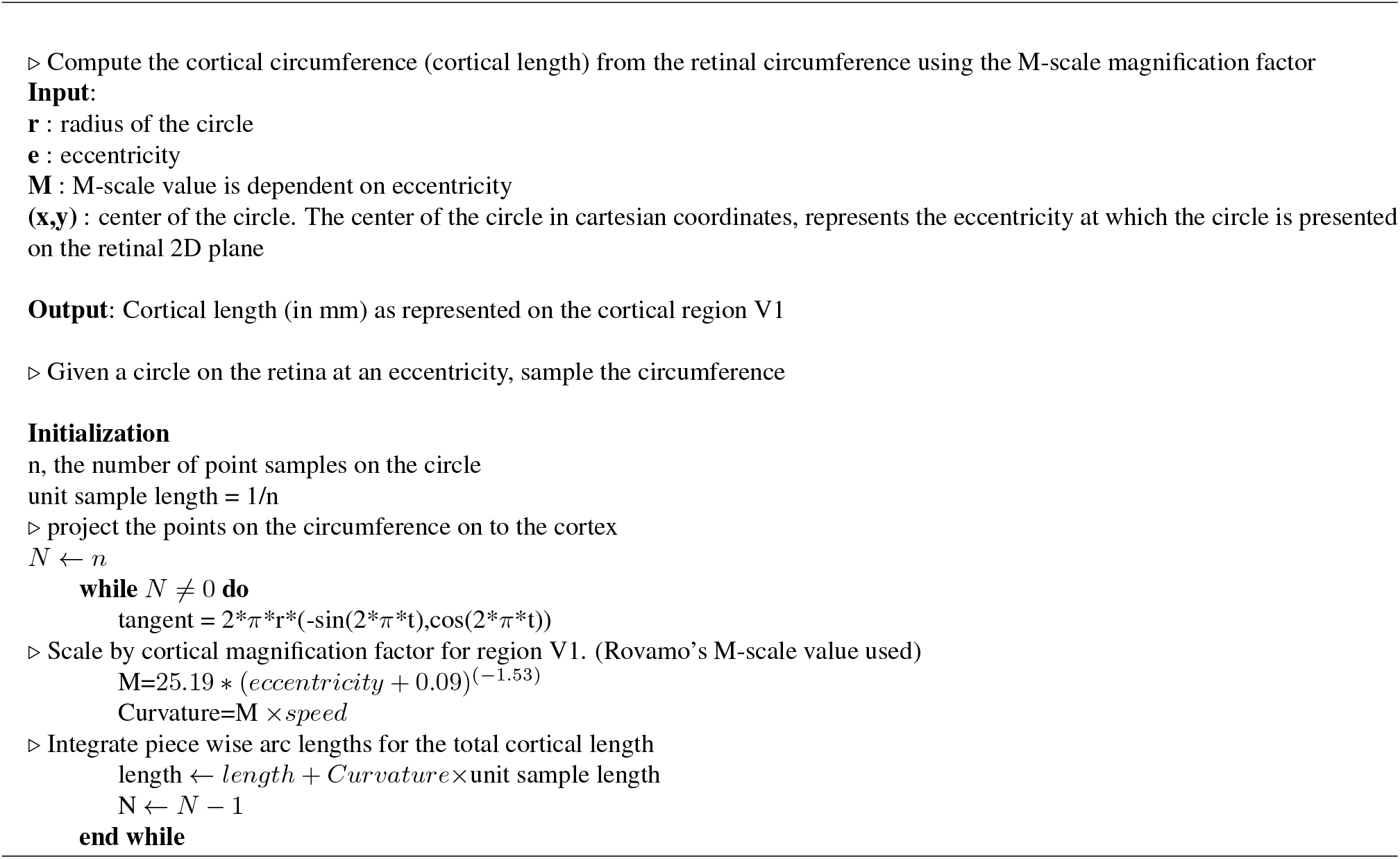

